# Telomere length set point regulation in human pluripotent stem cells critically depends on the shelterin protein TPP1

**DOI:** 10.1101/298778

**Authors:** John M. Boyle, Kelsey M. Hennick, Samuel G. Regalado, Jacob M. Vogan, Xiaozhu Zhang, Kathleen Collins, Dirk Hockemeyer

**Affiliations:** Department of Molecular and Cell Biology, University of California, Berkeley, Berkeley, CA 94720, USA

**Keywords:** Homeostasis, pluripotent stem cells, Telomere length, Telomere length set point, TPP1

## Abstract

Telomere maintenance is essential for the long-term proliferation of human pluripotent stem cells, while their telomere length set point determines the proliferative capacity of their differentiated progeny. The shelterin protein TPP1 is required for telomere stability and elongation, but its role in set point establishment remains elusive. Here, we characterize the contribution of TPP1 isoforms and residues outside the TEL patch, TPP1’s telomerase interaction domain, to telomere length control. We demonstrate that TPP1L, the longer minor isoform of TPP1, can partially compensate for loss of the more abundant shorter isoform, TPP1S. Both TPP1S knockout and complete TPP1 knockout cell lines (TPP1 KO) show telomere shortening. However, TPP1S KO cells are able to stabilize short telomeres while TPP1 KO cells do not and die. We compare these phenotypes with that of TPP1^L104A/L104A^ mutant cells that like the TPP1S KO have short stable telomeres. In contrast to TPP1S KO, TPP1^L104A/L104A^ cells respond to increased telomerase. However, TPP1^L104A/L104A^’s sensitivity to shelterin-mediated feedback is altered, revealing TPP1^L104A/L104A^ as a new type of shelterin mutant with aberrant set point regulation.

## Introduction

Telomere length maintenance in human stem cells is essential for their long-term proliferation and thus is linked to the renewal capacity of human cells and tissues (Aubert & Lansdorp, 2008). The enzyme telomerase catalyzes the addition of telomeric repeats to the chromosome end (Greider & Blackburn, 1985, Greider & Blackburn, 1989, Hemann, Hackett et al., 2000), thereby counteracting the terminal sequence loss by the end replication problem and nucleolytic degradation. In contrast to stem cells, most of their differentiated progeny downregulate telomerase resulting in progressive telomere shortening. This telomere shortening eventually leads to the induction of a DNA damage signal and cellular arrest or death (d’Adda di Fagagna, Reaper et al., 2003). This proliferation barrier, induced by progressive telomere erosion, functions as a strong tumor suppression mechanism (Shay, 2016). Human embryonic stem cells (hESCs) and induced pluripotent stem cells (iPSCs), collectively referred to as human pluripotent stem cells (hPSCs), resemble cells of the early human embryo that are telomerase positive and immortal (Takahashi, Tanabe et al., 2007, Thomson, Itskovitz-Eldor et al., 1998). hESCs maintain telomeres within a defined range, referred to as the telomere length set point, of approximately 9-12 kb (Rivera, Haggblom et al., 2016, Thomson et al., 1998). Analogously, iPSCs reset their telomeres to a similar length during the reprogramming process (Batista & Artandi, 2013, Batista, Pech et al., 2011). Additional evidence for a stem cell specific telomere length set point comes from experiments that revert TERT (Telomerase reverse transcriptase) loss of function in genetically engineered TERT deficient hESCs (Chiba, Johnson et al., 2015, Chiba, Vogan et al., 2017b). Restoration of endogenous TERT resets telomere length to the wild type set point. This argues that, in stem cells, a telomere length set point is a unique feature of telomere length control; it is distinct from merely counteracting telomere shortening and maintaining telomeres at homeostasis. In order to maintain homeostasis, a balance must exist between telomere sequence loss during DNA replication and *de novo* repeat addition. In order to establish homeostasis at a specific telomere length set point, stem cells must sense the total telomeric sequence present at a chromosome end, integrate this information with telomerase levels and mediate its access to the chromosome end. Based on experiments in model organisms and in cancer cells, it has been postulated that homeostasis is maintained by balancing telomeres between an extendable state and a non-extendable state (Teixeira, Arneric et al., 2004), which can be modulated depending on the presence of the shelterin complex at the telomere (Cristofari & Lingner, 2006, Hockemeyer & Collins, 2015, Hug & Lingner, 2006, van Steensel & de Lange, 1997). However, little is known about the mechanism by which shelterin regulates the telomere length set point. A defined telomere length set point is a specific feature of stem but not cancer cells. In cells of the same tumor type (Ceccarelli, Barthel et al., 2016, Hayward, Wilmott et al., 2017), and in subclones as well as second round subclones of cancer cell lines, telomere length can differ substantially (Bryan, Englezou et al., 1998).

Shelterin is comprised of six protein subunits (TRF1, TRF2, Rap1, TIN2, TPP1 and POT1) that are specifically recruited to the telomeric repeat array (de Lange, 2005, Schmutz & de Lange, 2016). This complex is required to protect the chromosome ends from an aberrant DNA damage response (DDR) (de Lange, 2010), and is required for telomere maintenance by telomerase. TRF1 and TRF2 bind to the double stranded telomeric repeats and have been shown to act as negative regulators of telomere length (Smogorzewska, van Steensel et al., 2000, van Steensel & de Lange, 1997). As it is the double stranded array that changes as telomeres shorten, current models suggest that telomere length is sensed by the binding of TRF1 and TRF2. TRF1 and TRF2, which also bind RAP1, are bridged by TIN2. TIN2 acts to recruit TPP1 (Houghtaling, Cuttonaro et al., 2004, Liu, Safari et al., 2004, Ye, Hockemeyer et al., 2004) and its binding partner, POT1. POT1 can bind single stranded telomeric DNA through its N-terminal OB-fold domains and controls telomerase access to the 3’-OH (reviewed in (Hockemeyer & Collins, 2015)).

TPP1 is essential in mice and for the proliferation of mouse keratinocytes, as well as mouse embryonic fibroblasts (MEFs) (Kibe, Osawa et al., 2010, Tejera, Stagno d’Alcontres et al., 2010). Conditional deletion of TPP1 elicits a DNA-damage response at telomeres, called telomere-dysfunction induced foci (TIFs) (Takai, Smogorzewska et al., 2003), cell cycle arrest due to p53 activation, and occasional chromosomal fusions, predominantly of sister chromatids. Overall, the mouse TPP1 loss of function phenotype closely resembles that of POT1 loss of function (Hockemeyer, Palm et al., 2007, Kibe et al., 2010, Kim, Li et al., 2017, Takai, Kibe et al., 2011). It is important to note, however, that analysis of the adrenocortical dysplasia (acd) mouse suggests that very little TPP1 is required for cellular and organismal viability (Hockemeyer et al., 2007, Keegan, Hutz et al., 2005). TPP1 is transcribed from the *acd* locus. The acd mouse is a mutant mouse strain that spontaneously arose. It suffers from reduced survival, reduced fecundity, poor growth, skin hyperpigmentation, and adrenal insufficiency. These phenotypes are caused by a homozygous mutation in intron 3 of TPP1 creating a cryptic splice donor that results in very low levels of TPP1 expression (approximately 2%) (Else, Theisen et al., 2007).

In its position bridging TIN2 and POT1, TPP1 can act to mediate telomere length signals coming from TRF1 and TRF2 and pass them to POT1 and telomerase. TPP1 is the only shelterin component shown to directly interact with telomerase through a motif referred to as the TEL patch. This interaction is essential for the recruitment of telomerase to telomeres (Hockemeyer & Collins, 2015, Nandakumar, Bell et al., 2012, Schmidt, Dalby et al., 2014, Sexton, Regalado et al., 2014, Wang, Podell et al., 2007, Xin, Liu et al., 2007, Zhong, Batista et al., 2012). hESCs harboring endogenous deletions of the acidic loop of TPP1’s TEL patch (TPP1^∆L/∆L^) show telomere shortening at rates that phenocopy those of telomerase knockout hESCs (Sexton et al., 2014). Complementation of TPP1^∆L/∆L^ hESCs with wild type TPP1 restored telomere length over time to the length of wild type hESCs. However, expression of a TPP1 variant containing a single amino acid substitution at position 104 (TPP1 L104A) outside of the TEL patch (Grill, Tesmer et al., 2018, Nandakumar et al., 2012), can rescue the long-term viability of TEL patch deficient cells but it fails to reset telomere length to wild type hESC levels (Sexton et al., 2014). The TPP1 mutant was originally described in an *in vitro* assay using purified TPP1 fragments missing the TIN2 binding domain (Grill et al., 2018, Nandakumar et al., 2012). This fragment was shown to have a defect in activating telomerase processivity however, in the context of the full length allele telomerase recruitment was shown to be unaffected (Grill et al., 2018).

To better dissect the role of TPP1 in telomere length set point control, we first characterized the two dominant isoforms of TPP1 present in cells by generating specific knockout cell lines for each isoform. We then contrasted these loss of isoform phenotypes with the phenotypes of cell lines containing endogenous homozygous TPP1 L104A mutations (TPP1^L104A/L104A^). This approach allowed us to demonstrate that TPP1 L104A is not a hypomorphic allele and does not lead to telomere deprotection, phenotypes that we find associated with the full TPP1 knockout and deletion of the most abundant isoform of TPP1. TPP1^L104A/L104A^ hESCs are capable of telomere end protection and homeostasis, which indicates that TPP1 L104 is competent to recruit and activate telomerase and tether POT1 to telomeres. However, TPP1^L104A/L104A^ hESCs show a pronounced defect in the stem cell specific telomere length set point and only maintain very short telomeres. The nature of the TPP1 defect caused by the L104A mutation was revealed by overexpressing well characterized alleles of telomeric binding proteins in TPP1^L104A/L104A^ hESCs. We show that TPP1 L104A presents a previously uncharacterized type of shelterin mutation that causes a severely blunted response to telomere length cues necessary to establish the hESC-specific telomere length set point.

## Results

### The human ACD (TPP1) locus expresses two isoforms in hESCs

To better understand the role of TPP1 in telomere length regulation in hESCs, we first characterized the TPP1 protein isoforms and their specific loss of function phenotypes. TPP1 has many annotated splice variants. However, recent polysome profiling and Transcript Isoforms in Polysomes sequencing (TrIP-seq) (Blair, Hockemeyer et al., 2017, Floor, Doudna et al., 2016) data suggests that hESCs express at least two isoforms of TPP1: a long isoform TPP1L (ENST00000393919), which starts translation at base pair 338 of the longest NCBI annotated transcript; and a second more abundant short isoform TPP1S (ENST00000620761), which is generated from a different transcript with a translational start site 258 bp downstream of TPP1L’s ATG (Fig 1A and EV1A). This downstream ATG encodes methionine 87 in the long transcript, hence, we refer to it as M87 where the first ATG in this transcript is referred to as M1 (Fig 1A). To address how each isoform individually contributes to telomere protection, length control and homeostasis, we mutated the respective ATG start codons of each isoform using CAS9-mediated genome editing (Jinek, Chylinski et al., 2012). Following editing we identified two clones with mono-allelic knock in of the repair template changing the short ATG to ATT (TPP1^M87I/+^) and one clone with mono-allelic knock in and disruption of the other allele leading to a premature stop (TPP1^M87I/−^). We refer to this latter clone as TPP1S KO. We also identified clones harboring either monoallelic and biallelic mutation at the first ATG (TPP1^∆M1/+^ and TPP1^∆M1/∆M1^). We refer to the biallelic disrupted cell line as TPP1L KO (Fig EV 1B, EV 1C). Both TPP1L and TPP1S KO cells remained pluripotent (Fig. 1B) and initially proliferated indistinguishably from unedited control clones and wild type cells. Mutation of the translational start of TPP1S resulted in the loss of a detectable protein product by Western blot. In contrast, mutation of the TPP1L ATG did not result in the depletion of a specific band (Fig 1C). This could be due to very low expression of TPP1L (Fig EV1A) being masked by a non-specific band at the expected size of TPP1L. An alternative explanation could be that the TPP1L ATG mutant cells initiate TPP1 translation on a downstream ATG at methionine 11; computational analysis of the of the TPP1L transcript with the M1 mutation by *ATG*^*pr*^ (http://atgpr.dbcls.jp/) shows an increased probability that M11 is used as a translational start site.

**Figure 1.**
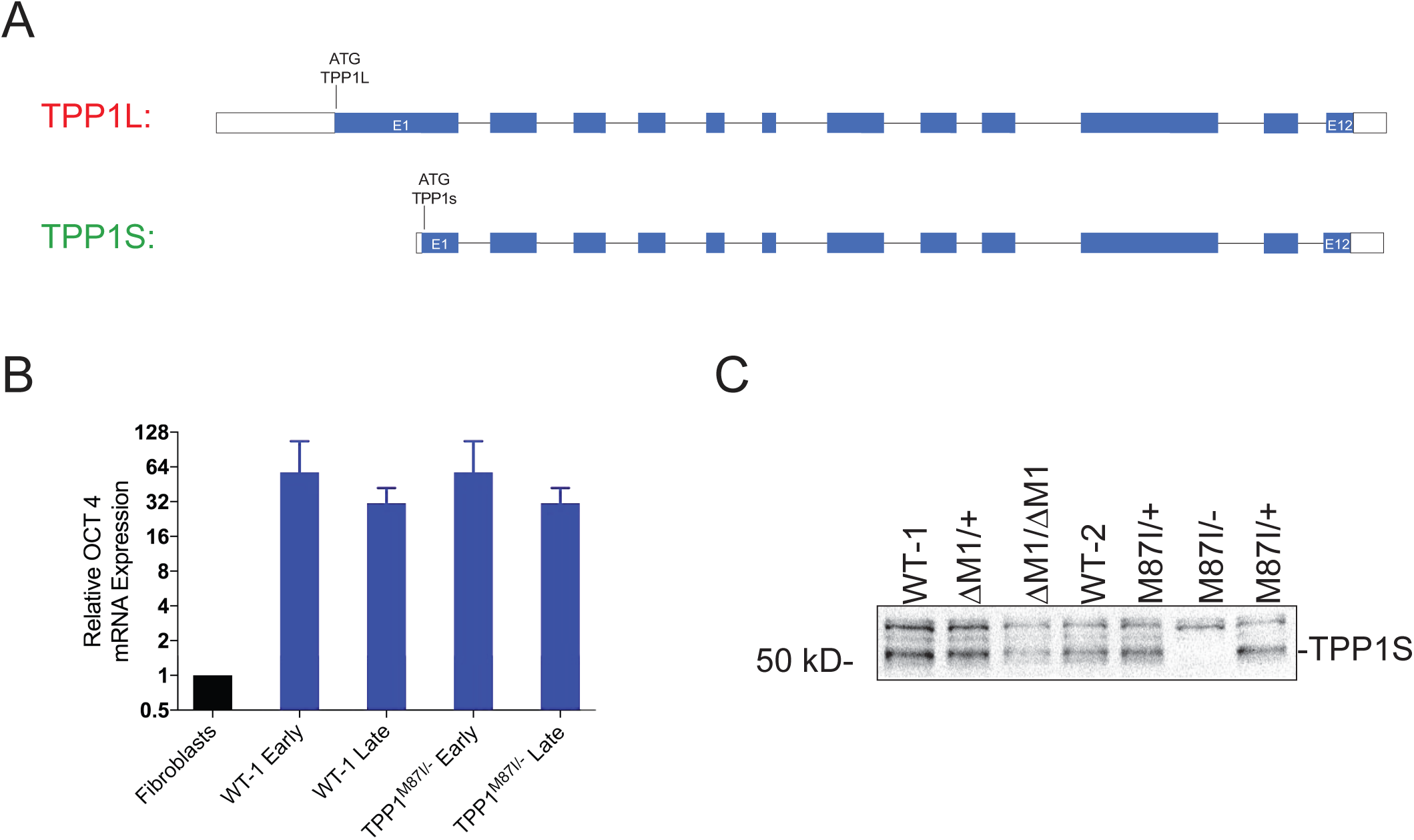
Targeted genome editing of the ACD locus. **A.** Schematic of the TPP1L, transcript ID ENST00000393919, and TPP1S transcript ID ENST00000620761. TPP1S is in frame with TPP1L and begins at the ATG 258 base pairs downstream from the start codon used by TPP1L. **B.** Relative OCT4 expression in early and late stage wild type and TPP1S KO cells compared to human fibroblasts (OCT4 negative). Each sample was normalized to GAPDH expression. Early samples were collected before noticeable proliferation defect on days 94, 113, and 120 following targeting. Late samples were collected after proliferation had stabilized on days 225, 240, and 261 following targeting. **C.** Western blot analysis of TPP1 gene products showing loss of TPP1S in TPP1^M87I/−^ as well as retention of all bands in WT, M1 homo and heterozygous deletions. Protein samples collected from cells 80 days following targeting.

### Loss of the predominant isoform TPP1S leads to telomere shortening and deprotection

To assess the proliferative capacity of the TPP1S and L knockout cell lines, we monitored telomere length and cellular proliferation (Fig 2A-C) over time. Initially, all cells grew at a rate indistinguishable from wild type. However, at approximately day 165 post targeting, TPP1S KO hESC cultures started to show a slower rate of proliferation, resulting in smaller stem cell colonies, significantly lower cell numbers, and increased number of dead cells detaching from the colonies and floating in the media (Fig 2C). This phenotype was highly reminiscent of late passage TERT^−/−^ and the telomerase recruitment mutant TPP1^∆L/∆L^ (Sexton et al., 2014). In contrast to TERT^−/−^ and TPP1^∆L/∆L^, TPP1S KO cultures continued to proliferate, but slower than wild type control cells. Following nine weeks of reduced proliferation, TPP1S KO cultures returned to a rate similar to wild type cells. Genotyping analysis of the surviving cell population ruled out potential cross contamination. TPP1S KO was also re-confirmed by western blot analysis in long-term culture after wild type proliferation rates had resumed (Fig 2D). TPP1L KO hESCs did proliferate continuously at the same rate as wild-type cells (Fig 2C).

**Figure 2.**
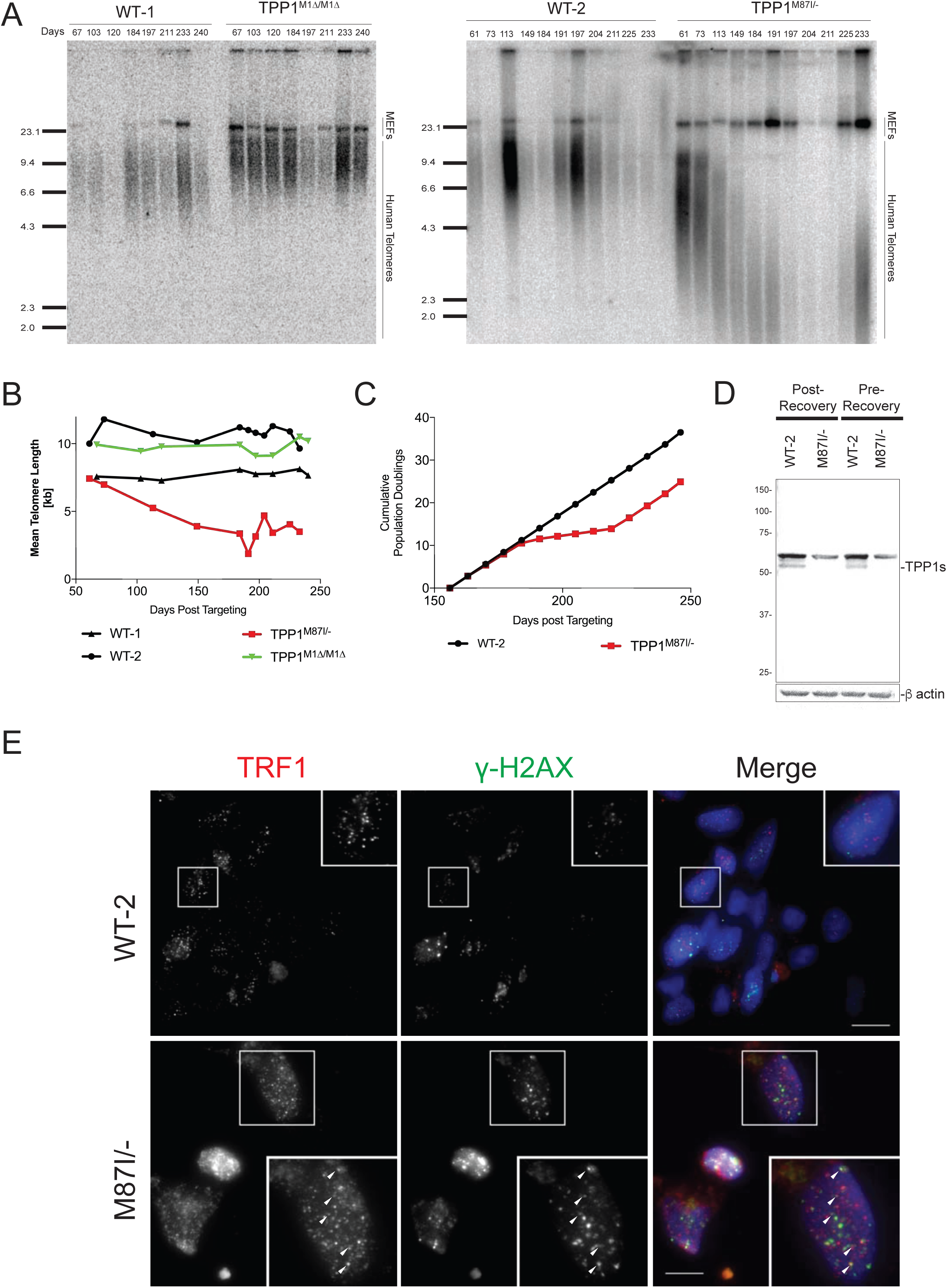
Effects of TPP1 isoform loss on telomere length and end protection. **A.** Telomere restriction fragment (TRF) analysis monitoring telomere length changes in wild type, TPP1L KO, and TPP1S KO cell lines over time. Numbers indicate days following targeting. **B.** Quantification of telomere length changes shown in A. **C.** Proliferation changes in TPP1S KO cell line compared to wild type. Changes in doubling time were calculated using the differences in split ratio used during passaging TPP1S KO cells versus WT-2 cells. Population doubling (PD) times were measured by taking the inverse of the fraction of cell passaged taken to the power of two. This number was added to the previous passages to give “cumulative PDs”. **D.** Western blot analysis confirming continued loss to TPP1S before and after the period of reduced proliferation. This indicated that proliferation recover was not the result of wild type contamination. Post-Recovery samples were collected on day 260 following targeting. Pre-Recovery samples were collected before noticeable proliferation defect was detectable on day 103 following targeting. **E.** Telomere-dysfunction Induce Foci (TIF) analysis of wild type and TPP1S KO cell lines. Arrows indicate colocalization of TRF1 and γ-H2Ax foci. Scale bars = 10μm. Sample were collected in parallel after recovery from proliferation defect on day 274 following targeting.

Next, we analyzed the telomere length changes in both TPP1L KO and TPP1S KO cells over time. This analysis revealed pronounced telomere shortening in TPP1S KO cells over the first 184 days. However, telomere shortening eventually slowed and mean telomere length stabilized at approximately 3.5 kb, and from there on telomere length remained stable. In contrast, telomeres in TPP1L KO and control cells did not shorten (Fig 2A, 2B, EV1C). Having established the differential function of the TPP1 isoforms in telomere length control, we next investigated their role in suppressing telomere DDR by quantifying the frequency of TIFs. While TIFs were absent from the wild type control and TPP1L KO cells, they are readily detectable in TPP1S KO cells (>37% of cells Fig 2E).

### Simultaneous disruption of both TPP1 isoforms leads to phenotypes different from the TPP1S KO

After observing the dramatic differences in telomere length and end protection between TPP1S and TPP1L KOs, we attempted to completely knockout TPP1. We made multiple attempts using different strategies to knockout TPP1 in wild type cells without success. We hypothesized that the failure to knock out TPP1 was the result of DNA damage checkpoint activation and p53-response (Kibe et al., 2010, Tejera et al., 2010). Therefore, we repeated the knockout in an hESC line with inactivated cell cycle and DNA damage checkpoints. We used a genetically engineered a hESC line with a biallelic deletion of exon two of CDKN2A, which inactivates both p14 and p16 (Chiba, Lorbeer et al., 2017a). In this cell line, we had also disrupted the *TERT* locus and complemented this loss by introducing a loxP flanked hTERT expression cassette into the *AAVS1* locus. Using this cell line, we were able to derive one single TPP1 knockout hESC clone that carried a homozygous integration of a PGK-PURO cassette (TPP1 ^E2-Puro/E2-Puro^), which is predicted to result in the complete loss of TPP1 activity (Fig 3A 3B, EV2). Telomeres in this clone rapidly shortened, resulting in cell death and eventual loss of all cells after approximately 87-120 days (Fig 3C). In an independent study of three clones from the parental cell line, hTERT loss resulted in cell death after 118, 131 and 145 days (Fig 3D). As expected, staining for γ-H2AX and TRF1 confirmed the presence of TIFs in the TPP1 ^E2-Puro/E2-Puro^ cells (Fig 3E, 3F). These data confirm that TPP1 is essential for long term stem cell viability.

**Figure 3.**
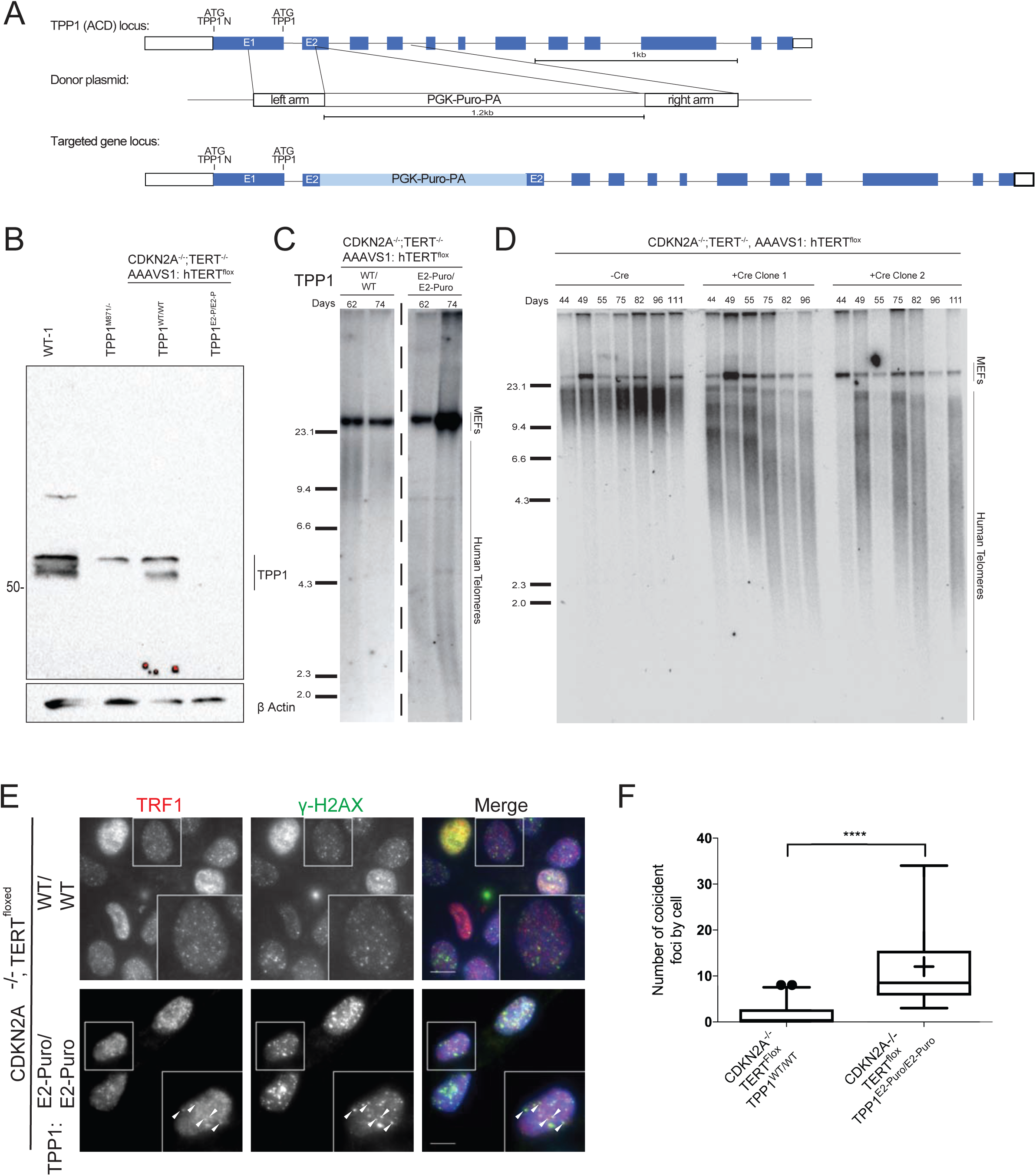
Analysis of complete loss of TPP1 in checkpoint deficient cells. **A.** Schematic depicting targeting strategy for knock in of a puromycin-poly A cassette into exon 2 of TPP1. **B.** Western blot analysis of TPP1 gene products in parental cell line and the cell line harboring the exon 2 insertion. WT-2 and TPP1S KO cells used as controls for gene product. M87I/− and control cells were collected 267 following targeting with TPP1S KO cassette. CDKN2A^−/−^ samples were collected 67 days following targeting with TPP1 exon 2 puromycin cassette. **C.** TRF analysis of TPP1 KO cell line. Numbers indicate days following electroporation. **D.** TRF analysis of parental CDKN2A^−/−^,TERT^−/−^, AAVS1:TERT^flox^ with and without Cre-mediated loop out of hTERT cassette. Numbers indicate days following transfection with either GFP mRNA or Cre mRNA. **E.** TIF analysis of CDKN2A^−/−^, TERT^−/−^, AAVS1:TERT^flox^ and CDKN2A^−/−^,TERT^−/−^, TPP1^E2-Puro/E2-Puro^, AAVS1:TERT^flox^ cell lines. Arrows indicate colocalization of TRF1 and γ-H2Ax foci. Scale bars = 10μm. Samples were collected 67 days following targeting with TPP1 exon 2 puromycin cassette. **F.** Quantification of coincident foci from CDKN2A^−/−^, TERT^−/−^, AAVS1:TERT^flox^ and CDKN2A^−/−^, TERT^−/−^, TPP1^E2-Puro/E2-Puro^, AAVS1:TERT^flox^ cell lines. Significant determined by Mann-Whitney test, n= 44 CDKN2A^−/−^,TERT^flox^, TPP1^WT/WT^ and 18 CDKN2A^−/−^,TERT^flox^, TPP1^E2-Puro/E2-Puro^ cells. Boxes indicate interquartile range, cross indicates median, “+” indicates mean and whiskers indicate 5%-95% range. P value is < 0.0001.

### TPP1 L104 is required for the full activity of telomerase at hESC telomeres

The telomere length set point of the TPP1S KO after long-term culture is reminiscent of the telomere length phenotype we previously identified when the TPP1 L104A allele was overexpressed in TPP1^∆L/∆L^ cells. TPP1 L104A was able to restore long term viability in TPP1^∆L/∆L^ but resulted in uncharacteristically short telomeres for stem cells (Sexton et al., 2014). These complementation assays did not resolve the question of whether TPP1 L104A is a hypomorphic loss of function allele or acts as a neomorphic allele when expressed endogenously. To fully characterize this mutant and to determine whether TPP1 ^L104A/L104A^ will act in a similar manner as TPP1S KO, we edited the endogenous TPP1 locus changing leucine 104 to alanine (Fig 4 and EV3A-C). Telomere length showed that the homozygous TPP1^L104A/L104A^ substitution at the endogenous TPP1 locus results in the gradual loss of telomeric signal (Fig 4A, 4B, EV 3C, EV 3D). This loss continued for over 130 days following editing and finally stabilized at approximately 3.5 kb, similar to the length where TPP1S KO cells stabilized.

**Figure 4.**
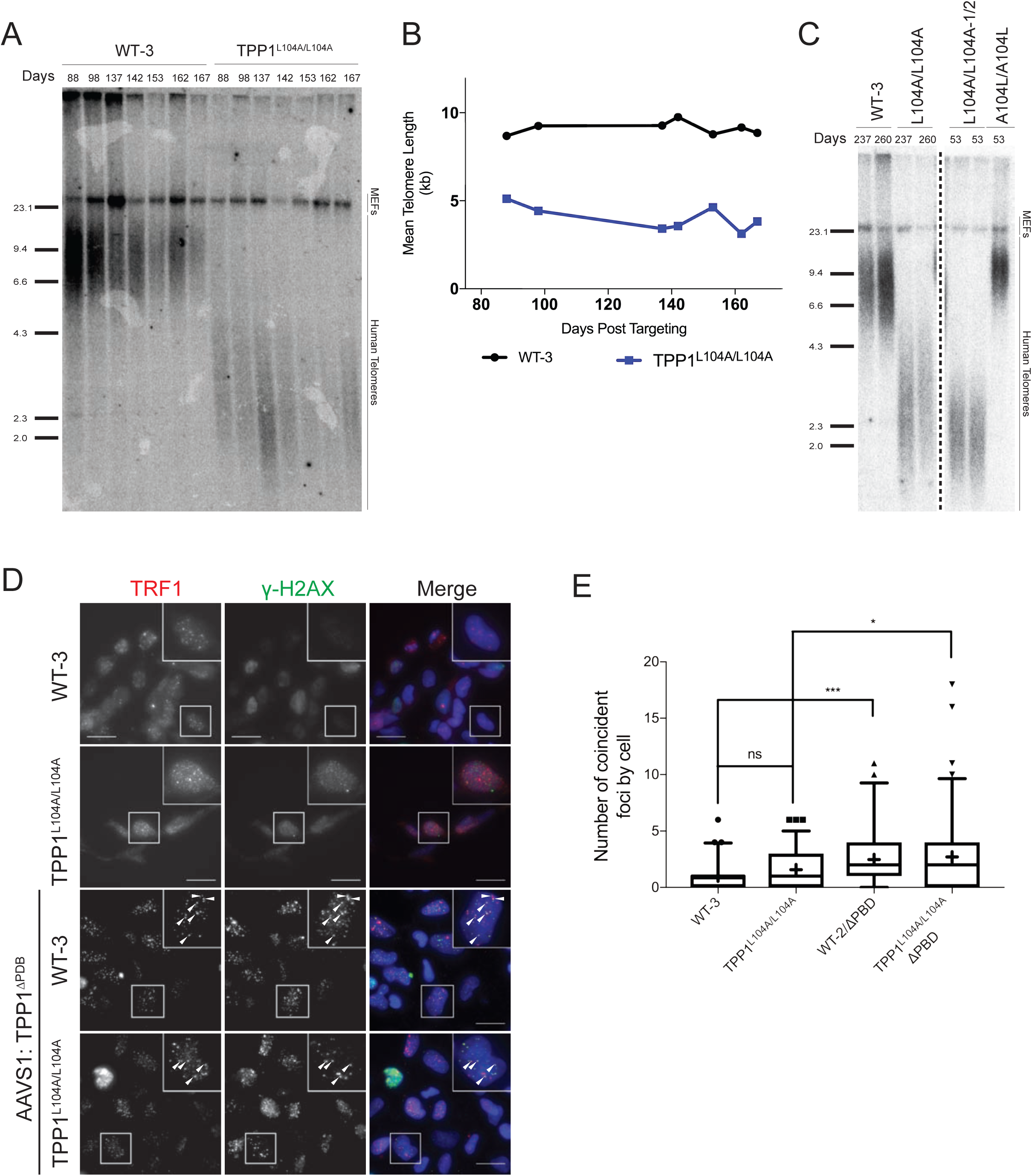
Effects of TPP1^L104A/L104A^ mutation on telomere length and end protection. **A.** TRF analysis monitoring telomere length changes in wild type and TPP1^L104A/L104A^ cell lines over time. Numbers indicate days following targeting. **B.** Quantification of telomere length changes shown in A. **C.** TRF analysis of telomere length changes following reversion of the TPP1 locus to wild type, parental mutants, and repaired cell lines. Two A104L/A104L clones were isolated and analyzed. Numbers to the left of the dashed line indicate days following targeting of original L104A mutant. Numbers to the right of the dashed line indicate days following targeting of the A104L repair. Sample from days 260-left and 53-right were collected on the same day. All samples were run on the same gel, dashed line indicates collapsed image. **E.** TIF analysis of wild type and TPP1^L104A/L104A^ cell lines. TPP1^∆PDB^ overexpression cell lines were included as a positive control. Cells were fixed 26 days following targeting the TPP1^∆PDB^ expression cassette into the *AAVS1* locus. TPP1^∆PDB^ cell lines were generated from parental cell lines that were originally targeted over 200 days prior and had stabilized their telomeres. Arrows indicate colocalization of TRF1 and γ-H2Ax foci. Scale bars = 10μm. **E.** Quantification of TIFs CDKN2A^−/−^, TERT^−/−^, AAVS1:TERT^flox^ and CDKN2A^−/−^,TERT^−/−^, TPP1^E2-Puro/E2-Puro^, AAVS1:TERT^flox^ cell lines. Significance determined by Mann-Whitney test, n= 60 WT-3, 91 TPP1^L104A/L104A^, n= 54 WT-2; AAVS1: TPP1^∆PBD^, and n= 86 TPP1^L104A/L104A^; AAVS1: TPP1^∆PBD^ cells. Boxes indicate interquartile range, cross indicates median, “+” indicates mean and whiskers indicate 5%-95% range. “****” indicates a P value < 0.0001 and “*” indicates a P value < 0.05.

In order to confirm the L104A allele is not a hypomorph, we used overexpression of TPP1 alleles from the *AAVS1* locus to further characterize the genetic basis of the TPP1^L104A/L104A^ mutation in hESCs. Overexpression of TPP1 L104A and TPP1∆L failed to rescue the telomere length phenotype seen in TPP1^L104A/L104A^ mutant cells (Fig EV3D-F), whereas wild type TPP1S restored telomere length to the hESC telomere length set point. Furthermore, we noticed that residue 104 is located adjacent to lysine 233 in the folded protein (Fig EV4A). This lysine has been shown to be a ubiquitination target required for TPP1 stabilization in mice (Rai, Li et al., 2011). However, when we overexpressed TPP1S, TPP1S K232A, TPP1S K233A, or TPP1S KK232/3AA in TPP1^∆L/∆L^ cells, or TPP1S^K233A^ in the TPP1^L104A/L104A^ cells (Fig EV4B), telomeres were restored to their typical length. Therefore, the TPP1^L104A/L104A^ phenotype is distinct from TPP1S variants that cannot be ubiquitinated at lysine 232 and/or 233. Importantly, we confirmed the specificity of the TPP1^L104A/L104A^ telomere length phenotype by genetically reverting the defect. The repair of TPP1^L104A/L104A^ to TPP1^A104L/A104L^ resulted in the rapid reestablishment of the telomere length to the hESC specific set point, confirming the specificity of the TPP1^L104A/L104A^ phenotype (Fig. 4C). These results show the L104A allele is a novel telomere length set point mutant that is not caused by generating a hypomorphic protein.

### Telomerase levels increase over time in TPP1S knockout cells but not TPP1 L104A cells

Both hESC lines, TPP1^L104A/L104A^ and TPP1S KO, have shortening telomeres that eventually reach homeostasis at approximately 3.5 kb. Thus, in the beginning there is a net loss of telomeric DNA, but over time telomere repeat addition increases to a level that is sufficient to establish homeostasis. Next, we investigated if changes in telomerase activity are the basis for the restoration of homeostasis. Using the PCR-based telomeric repeat amplification protocol (TRAP) to assay telomerase activity of early and late TPP1S KO cells revealed an increase in activity at the time when TPP1S knockout showed reduced proliferation. Telomerase activity remained elevated in all later time points (Fig 5A). This suggests that TPP1S KO cells compensate for the reduced TPP1 function by increasing telomerase levels (Fig 5A). In contrast, TPP1^L104A/L104A^ stabilize telomeres without changing the overall telomerase levels, suggesting a different mechanism by which telomere length stabilizes (Fig 5B).

**Figure 5.**
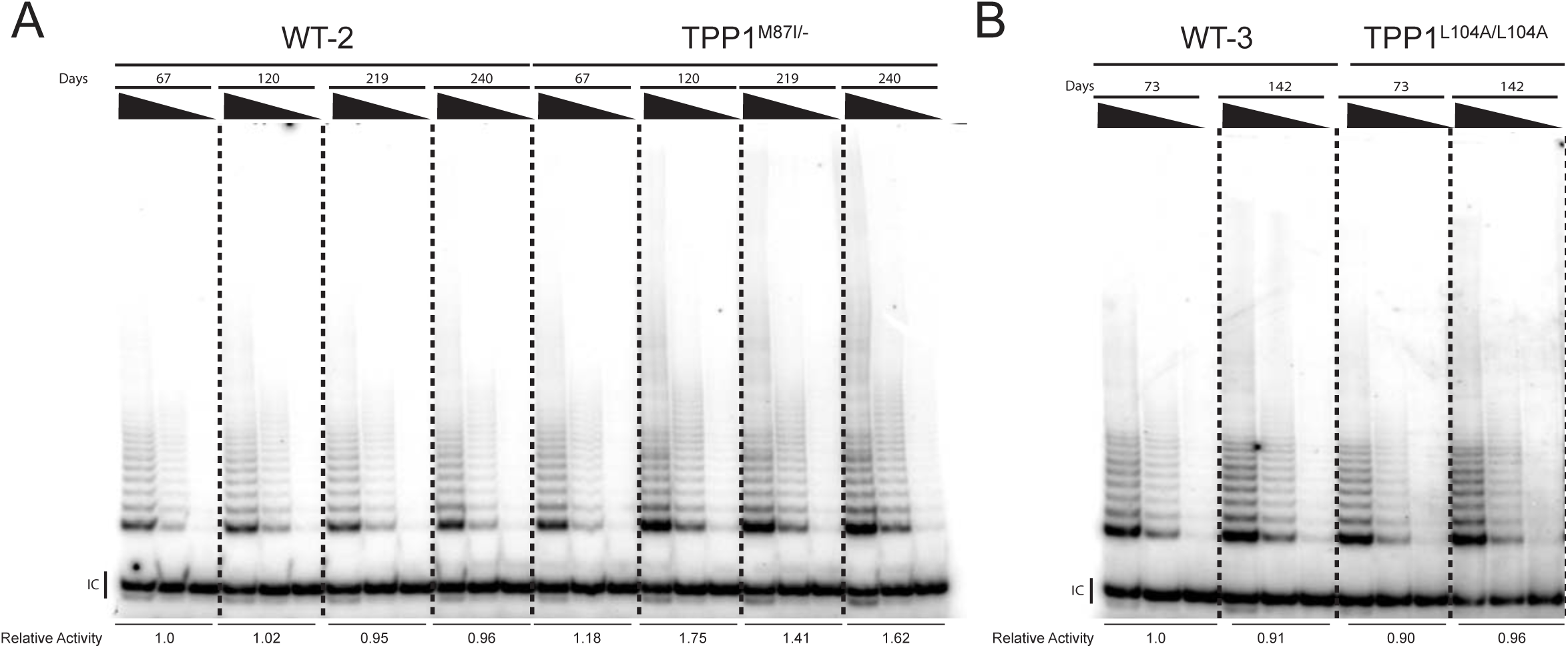
Telomerase activity changes in TPP1S KO cells, not TPP1^L104A^ cells. **A.** Analysis of telomerase activity by Telomere repeat addition assay (TRAP) in wild type, and TPP1S KO cell lines. Relative activity was determined using ImageJ to calculated total band intensity in first lane of each time point. Increased activity is evidenced after day 120 in the TPP1S KO cells. Numbers above the radiogram indicate days following targeting M87 generating the TPPS KO. **B.** TRAP analysis as in (A) but of wild type and TPP1^L104A/L104A^. Relative activity was determined as in (A). Numbers above the radiogram indicate days following targeting L104A.

### Telomeres are deprotected in TPP1S knockout, but not homozygous TPP1 L104A, hESCs

Since the new telomere set points of both L104A and TPP1S KO cells were approximately the same, we asked whether TPP1^L104A/L104A^ has deprotected telomeres, similar to the TPP1S KO cells. To answer this question, we assessed telomere protection by quantifying TIFs in TPP1^L104A/L104A^ and TPP1S KO cells. TPP1^L104A/L104A^ cells only showed wild type levels of TIF positive cells (Fig 4D, 4E) while TPP1S KO cells had high level of TIFs (Fig 2E). We overexpressed a TPP1 variant missing the POT1-binding domain (TPP1^∆PBD^) from the *AAVS1* locus to act as a positive control for TIF positive cells (Fig 4E). Expression of this TPP1 variant resulted in an increase of TIF positive cells in wild type and TPP1^L104A/L104A^ cells. These results indicate that TPP1^L104A/L104A^ is sufficient to protect short telomeres from a DDR. While telomere length is consistent between TPP1S KO and L104A cells, only the TPP1S KO showed telomere deprotection.

### TPP1 L104 is a novel threshold mutant of telomere length control

Investigating the difference between the TPP1 L104A mutant and the TPP1S KO further, we then asked if both mutants are responsive to manipulations known to promote telomere elongation. Overexpression of either POT1∆OB—an allele of POT1 that lacks the first OB-fold required for DNA binding (Loayza & de Lange, 2003)—or simultaneous expression of hTERT and hTR from the *AAVS1* locus led to rapid telomere elongation in wild type cells (Fig 6A) (Chiba et al., 2015, Hockemeyer, Soldner et al., 2009, Hockemeyer, Wang et al., 2011). Although TPP1S KO cells showed telomere elongation after either POT1∆OB or telomerase expression, the response was severely muted. Overexpressing the same alleles in TPP1^L104A/L104A^ cells led to a more complex outcome. Overexpression of telomerase leads to fast telomere elongation, driving telomeres beyond the set point of wild type cells. This was different from the expression of POT1∆OB, where telomeres slowly elongated and nearly reached the wild type telomere length set point (Fig 6A). This failure of TPP1^L104A/L104A^ cells to appropriately respond to changes in shelterin becomes even more pronounced when analyzing the telomere elongation following overexpression of a dominant negative form of TRF1 missing the acidic and Myb domains (TRF1^∆A∆M^) (Fig. 6B, EV5). These data suggest that the L104A allele is more responsive to the downstream effectors of telomere length, POT1 and telomerase, and less responsive to the upstream effectors, TRF1 and TRF1^∆A∆M^.

**Figure 6.**
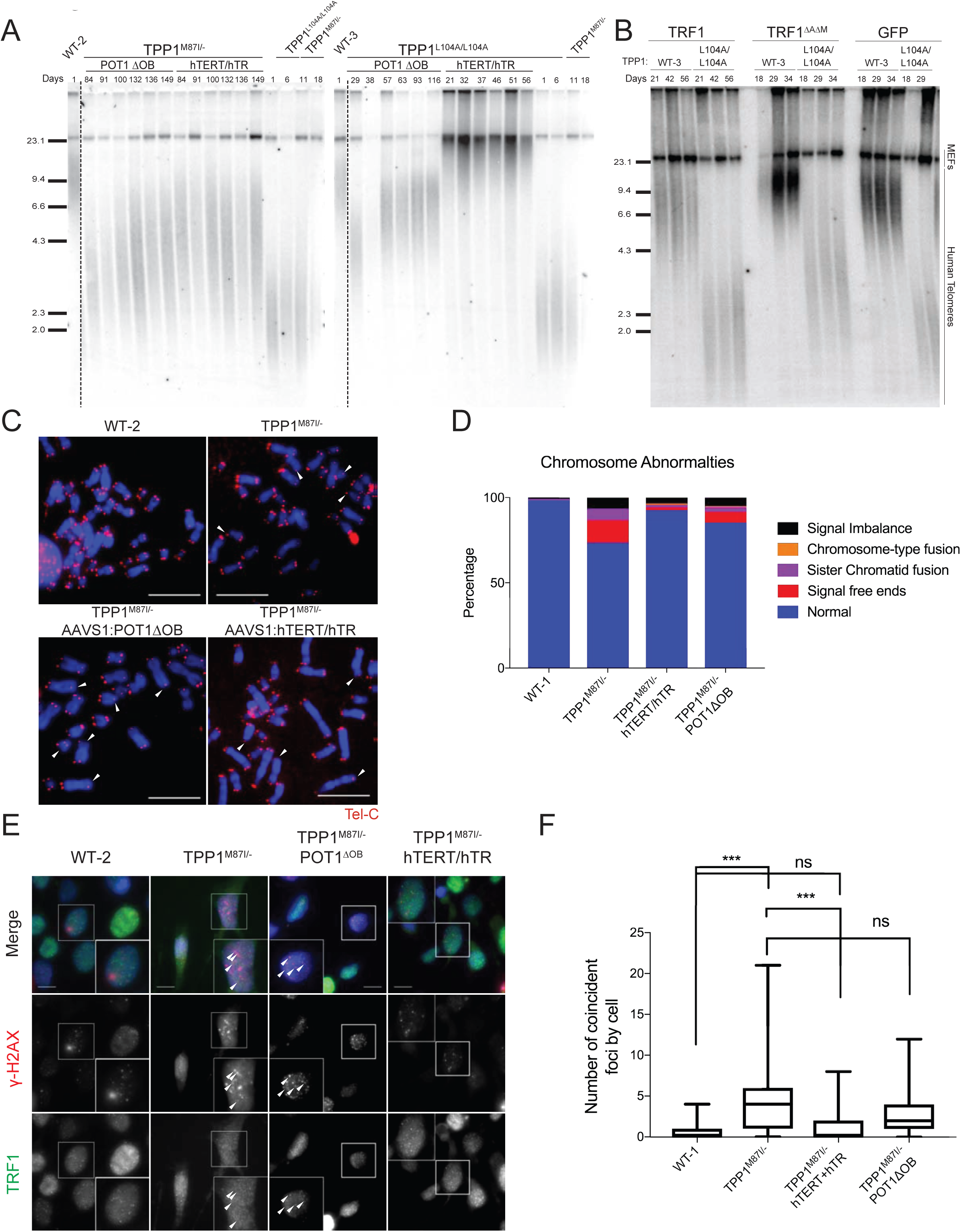
Response of TPP1S KO and TPP1^L104A/L104A^ cell lines to telomere length signals. **A.** TRF analysis of telomere length changes in wild type, TPP1S KO, and TPP1^L104A/L104A^ cell lines following overexpression of either POT1∆OB or hTERT/hTR, downstream regulators of telomere length. Numbers indicate days following targeting of the *AAVS1* locus with either a POT1∆OB or hTERT/hTR overexpression cassette. TPP1^M87I/−^ parental cell line as well as WT-2 control are synchronized with TPP1^M87I/−^ overexpression cells lines; day 1 is 164 days post original KO targeting. TPP1^L104A/L104A^ parental cell line as well as WT-3 control line are synchronized with TPP1^L104A/L104A^ overexpression cells lines; day 1 is 136 days post original L104A targeting. All samples were run on two gels, dashed line indicates collapsed image. **B.** TRF analysis of telomere length changes in wild type and TPP1^L104A/L104A^ cell lines following overexpression of either TRF1, TRF1^∆A∆M^, a dominant negative allele of TRF1, or GFP control. TRF1 is thought to act upstream of TPP1 in telomere length regulation. Numbers indicate days following targeting the *AAVS1* locus with TRF1^∆A∆M^ or GFP a cassette overexpression cassette. **C.** Metaphase spreads of wild type, TPP1S KO cell line with and without over expression of either POT1∆OB or hTERT/hTR. Metaphases were fixed 240 days following targeting for WT-2 and TPP1^M87I/−^ and day 89 following targeting of the *AAVS1* locus for overexpression cell lines. Metaphases were processed in parallel. Scale bars = 10μm. **D.** Quantification of metaphase spreads represented in (C). 302 WT-1, n= 119 TPP1^M87I/−^, n= 716 TPP1^M87I/−^, AAVS1: POT1∆OB, and n= 462 TPP1^M87I/−^, AAVS1: hTERT/hTR chromosome ends. Cells were fixed 96 days following targeting of the *AAVS1* locus. **E.** TIF analysis of WT-2; TPP1^M87I/−^; TPP1^M87I/−^, AAVS1:POT1∆OB; and TPP1^M87I/−^, AAVS1:hTERT/hTR cell lines. Arrows indicate colocalization of TRF1 and γ-H2Ax foci. Scale bars = 10μm. Samples were collected 105 day after targeting the *AAVS1* locus. **F.** Quantification of coincident foci represented in (E). Significant determined by Mann-Whitney test, n= 65 WT-1; n= 29 TPP1^M87I/−^; n= 60 TPP1^M87I/−^, AAVS1: POT1∆OB; and n= 35 TPP1^M87I/−^, AAVS1: hTERT/hTR cells. Boxes indicate interquartile range, cross indicates median, “+” indicates mean and whiskers indicate 5%-95% range. “***” indicates a P value < 0.001.

As TPP1S KO cells showed a severely muted response to telomere lengthening cues, we next investigated to what extent these manipulations could restore telomere protection. Analyzing metaphase spreads of TPP1S KO cells revealed a significant number of chromosomal and telomere specific defects in TPP1S KO cells, including telomere signal loss, unbalanced telomere signals, and sister chromatid fusions. In contrast, we did not see a significant number of aberrant chromosomes in TPP1 ^L104A/L104A^ cells. Interestingly, while expression of either POT1∆OB or hTERT/hTR in TPP1S KO cells has very little effect on telomere length, their overexpression is capable of partially suppressing the metaphase phenotypes seen in TPP1S KO cells. (Fig 6C, 6D). Telomerase-mediated suppression was more pronounced when compared to the POT1∆OB-mediated suppression, yet expression of POT1∆OB significantly increased telomere protection. We also examined TIFs in the TPP1S KO cells and found that telomerase expression significantly suppressed the TIF phenotype in TPP1S KO cells, while expression of POT1∆OB resulted did not result in a significant reduction (Fig 6E, 6F).

## Discussion

### TPP1 has a critical function in establishing the telomere length set point in stem cells

Telomere length set point is determined by the interplay of two inputs: a repressive signal coming from TRF1 and TRF2 sensing the length of the double stranded telomeric DNA, and telomerase levels (Hockemeyer & Collins, 2015). It is on TPP1 that these two signals converge and are integrated to establish a telomere length set point and maintain homeostasis. The only requirement on telomere length in cancer cells is to maintain telomeres over time above a critical length. However, most cancer cells and cancer cell lines lack the genetic and epigenetic stability to maintain the levels of telomerase activity and TRF1/TRF2 signaling to allow a fixed telomere length set point (Bryan et al., 1998). In contrast, stable telomerase activity and TRF1/TRF2 signaling is a hallmark of hPSCs. This stability allows stem cells to maintain telomeres long enough to allow for the full development of an organism, but short enough for telomere shortening in differentiated progeny to function as a tumor suppression mechanism (Shay, 2016). Using this system, we have analyzed the contributions of TPP1 isoforms to each of these roles and identified a specific mutation in TPP1 that explicitly alters the stem cell telomere length set point.

### Stem cells express at least two TPP1 isoforms

In our loss of function studies, we have identified specific roles for the different isoforms. Most overexpression studies to date evaluated TPP1 function only in the context of TPP1S but did not address if the longer isoform may contribute to the cellular function of TPP1. Using TrIP-seq data (Blair et al., 2017) and functional assays, we provide evidence for the expression of the longer TPP1 isoform, TPP1L. Isoform-specific knockouts of TPP1S and TPP1L show that the less abundant isoform is able to partially compensate for the loss of TPP1S. Cells lacking TPP1S are long-term viable while cells disrupted in both TPP1S and TPP1L are not. This argues that the expression of TPP1L is sufficient for the immortal phenotype of stem cells. Expression of TPP1L alone, while capable of maintaining viability, is not sufficient for telomere protection and genome integrity, and hESCs with only TPP1L have telomeres that are maintained at a dramatically shorter length set point than typical for stem cells. However, our experiments cannot rule out the possibility of other TPP1 species compensating for the loss of TPP1S.

### TPP1S deficiency generally impairs TPP1 functions while the TPP1 L104A mutation results in a specific length set point defect

Maintaining telomere length homeostasis is only one role of TPP1. We further investigated the role TPP1 has in establishment of telomere length set point and chromosome end protection mediated through POT1 recruitment. The phenotype of TPP1S KO cells is reminiscent of that seen in the TPP1 L104A complementation of TPP1^∆L/∆L^ mutant cell lines. This led us to further investigate the similarities and differences between TPP1^L104A/L104A^ and TPP1 KO cell lines. The presence of TIFs and abnormal chromosome ends in TPP1S KO cells revealed that TPP1S is required to properly mediate end protection. A similar defect in telomere protection is not present in the L104A mutant, indicating that a TPP1S protein, harboring an L104A substitution, can properly interact with TIN2 and POT1.

During a period of low proliferation and increased cell death in TPP1S KO cells, telomerase activity increases. This increase in activity suggests that cells lacking TPP1S rely on changes in telomerase activity to restore telomere length homeostasis. In contrast, TPP1^L104A/L104A^ cells are able to maintain telomeres without a commensurate increase in telomerase activity.

The increase of telomerase activity, seen in TPP1S KO, may lead to the saturation of available TPP1. If true, we would predict that TPP1 is limited in these cells and the addition of more telomerase or other telomere lengthening cues would have no effect on telomere length. This hypothesis was confirmed; in the TPP1S KO cells, we report only a very small increase in telomere length in response to both of these cues. However, in the TPP1^L104A/L104A^ cells, we saw pronounced increase in telomere length following both hTERT/hTR and POT1∆OB overexpression. These data taken together demonstrate that loss of TPP1S results in hypomorphic TPP1 in all measured responses. On the other hand, L104A only shows defects in one known role of TPP1, establishment of telomere length set point.

### A model for how TPP1 coordinates the interconversion of telomeres from a telomerase extendable to non-extendable state

Current models of telomere length sensing invoke a still unresolved “counting” mechanism where TRF1 and TRF2 act to “count off” double stranded telomeric repeats. This signal is then passed to TPP1, which is then transduced to both telomerase and POT1 (Loayza & de Lange, 2003). The strength of this signal causes a change between extendable and non-extendable telomeric states (Cristofari & Lingner, 2006, Hug & Lingner, 2006, Smogorzewska & de Lange, 2004, Teixeira et al., 2004). As telomeres shorten, this balance shifts to a more extendable state allowing telomerase access to the 3’-OH and restoring the typical telomere length.

Our analysis of the TPP1 L104A mutant shows that TPP1 is an integral shelterin member responsible for balancing between a non-extendable and extendable state based on the upstream telomere length signal coming from TRF1 and TRF2. TRF1 and TRF2 act as negative regulators of telomere length (Smogorzewska et al., 2000, van Steensel & de Lange, 1997), indicating that they regulate telomere length homeostasis by sending a repressive signal. We propose a model for the L104A mutation in which the ability of TPP1 to sense the signal from TRF1 and TRF2 is more sensitive. When TPP1 L104A was knocked into wild type cells, the repressive signal from TRF1 and TRF2 is sensed very strongly and the equilibrium is pushed heavily toward non-extendable. As telomeres shorten, the repressive signal diminishes, and the equilibrium slowly shifts to a more extendable state. Finally, when the signal is low enough to counteract the hyper-sensitivity of TPP1^L104A/L104A^, we see the establishment of a new telomere length set point. At this new set point, the output from TPP1^L104A/L104A^ is equivalent to the output from wild type TPP1 at hESC typical set point. This model is consistent with TPP1^L104A/L104A^ cells being able to respond to hTERT/hTR or POT1∆OB signaling, but not to the same extent as wild type cells. Since POT1 and telomerase act downstream of TPP1, their overexpression can partially overcome the more repressive nature of the L104A mutant. The most severe failure to elongate telomeres was seen in TPP1 ^L104A/L104A^ cells overexpression of a dominant negative form of TRF1, suggesting that L104A is epistatic to TRF1. These findings establish the L104A mutation as a length threshold hypermorph, which means the main feature of the L104A mutant is an altered threshold at which it allows telomerase action at the chromosome ends (Fig 7).

**Figure 7.**
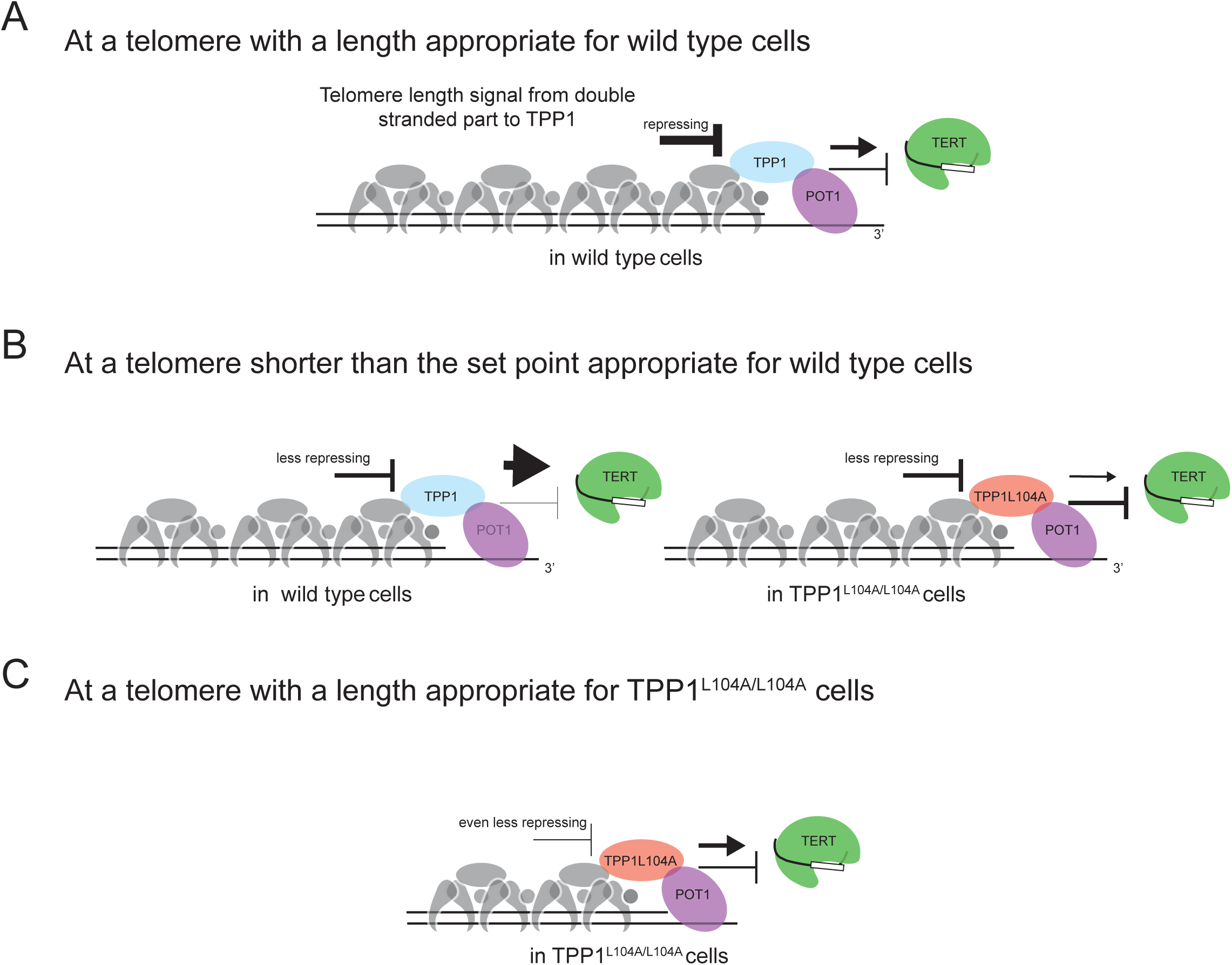
Model of L104-mediated telomere length set point regulation. **A.** At typical telomere length in hPSC, TRF1 and TRF2 produce a repressive signal that acts on TPP1. This signal is integrated with telomerase activity and mediates access to the 3’-OH via POT1. **B.** As telomeres shorten this signal becomes less repressing. In cells with wild type TPP1, the decrease in repression results in TPP1 increasing recruitment and activation of telomerase and reduces sequestration of the 3’-OH. TPP1^L104A^ is more sensitive to the signal coming from the upstream shelterin components therefore in cells expressing only TPP1^L104A^, telomeres will continue to shorten. **C.** As telomeres in TPP1^L104A/L104A^ hESCs reach approximately 3.5 kb, the signal coming from TRF1 and TRF2 is reduced enough to elicit an output from TPP1^L104A^ that is identical to wild type TPP1 at typical stem cell telomere length. This establishes the new telomere length set point.

## Materials and Methods

### hESC culture and genome editing

Genome editing experiments were performed in WIBR#3 hESCs (Lengner, Gimelbrant et al., 2010), NIH stem cell registry # 0079. Cell culture was carried out as previously described (Soldner, Hockemeyer et al., 2009). For CDKN2A exon2 deletion, the region flanked by accattctgttctctctggc and cgcggaaggtccctcaggtg as sgRNA target sites was removed and genotyped as previously reported (Chiba et al., 2017a). For knockout and L104A knock in experiments, cells were co-electroporated with targeting and GFP plasmids, cells were single cell sorted by florescence-assisted cell sorting and single cell derived colonies were isolated, replicate plates were made, and genomic DNA was isolated for confirmation of genotypes. Targeting was confirmed by Sanger sequencing and Southern blot analysis using a probe amplified from genomic DNA using Fw:ggggaaatgatgttggcttagaatcct and Re:caatgaagtccttcgtcttggtca as primers. Southern blot analysis was performed as previously described (Hockemeyer et al., 2009, Hockemeyer et al., 2011). *AAVS1* targeting was performed as previously described (Sexton et al., 2014). Targeting construct contained cDNA expressing TPP1S-3xF, TPP1^∆PBD^-F, myc-POT1∆OB, F-hTERT/hTR, F-TRF1, or F-TRM1^∆A∆M^, respectively.

### Telomeric repeat amplification protocol

PCR-based telomeric repeat amplification protocol (TRAP) was performed as previously described using TS (AATCCGTCGAGCAGAGTT) and ACX (GCGCGGCTTACCCTTACCCTTACCCTAACC) for amplification of telomeric repeats, and TSNT (AATCCGTCGAGCAGAGTTAAAAGGCCGAGAAGCGAT) and NT (ATCGCTTCTCGGCCTTTT) as an internal control (Kim, Piatyszek et al., 1994).

Cell extracts were generated using Hypotonic Lysis buffer (HLB) (20mM HEPES, 2mM MgCl_2_, 0.2mM EGTA, 10% Glycerol, 1mM DTT, 0.1mM PMSF) supplemented with 0.5% CHAPS. Samples were normalized using the Bradford assay (Bio-Rad). Serial dilutions, 200 ng, 40 ng, and 5 ng, of whole cell lysate were used to assay telomerase activity. The TRAP products were resolved on 10% polyacrylamide /1xTAE gels. Dried gels were visualized by phosphor imaging.

### Immunoblotting

Protein samples were extracted in RIPA buffer, sonicated to liberate chromatin bound proteins, normalized for protein concentration, and mixed with Laemmli buffer. After heating to 95°C for 5 minutes, samples were resolved by SDS-PAGE. Proteins were then transferred to nitrocellulose membrane and subsequently incubated with rabbit α-TPP1 (1:1000, A303-069A, [Bethyl]) and mouse α-Actin (1:20000[Santa Cruz]) in 5% nonfat milk [Carnation] in TBS-T buffer (150 mM NaCl, 50 mM Tris pH 7.5, 0.1% Tween 20) overnight at 4°C. The membrane was washed in TBS-T and incubated with goat α –mouse/rabbit HRP (1:5,000, [Bio-Rad]) in 5% nonfat milk in TBS-T for 1 hr. at room temperature. After extensive washing with TBS-T, the membrane was visualized using ECL.

### Detection of telomere length

Genomic DNA was prepared as described previously (Hockemeyer, Sfeir et al., 2005). Genomic DNA was digested with MboI, AluI and RNaseA overnight at 37°C. The resulting DNA was normalized and 2µg of DNA was run on 0.75% agarose [Seakem ME Agarose, Lonza], dried under vacuum for 2h at 50°C, denatured in 0.5 M NaOH, 1.5 M NaCl for 30 mins, shaking at 25°C, neutralized with 1 M Tris pH 6.0, 2.5 M NaCl shaking at 25°C, 2x for 15 min. Then the gel was pre-hybridized in Church’s buffer (1% BSA, 1 mM EDTA, 0.5M NaP0_4_ pH 7.2, 7% SDS) for 1h at 55°C before adding a ^32^P-end-labeled (C_3_TG_2_)_3_ telomeric probe. The gel was washed 3x 30 min in 4x SSC at 50°C and 1x 30 min in 4x SSC + 0.1% SDS at 25°C before exposing on a phosphor imager screen.

### qRT-PCR

RNA was extracted with TRIzol [Lifetech] and treated with DNaseI [NEB]. 600ng RNA were converted to cDNA with iScript Reverse Transcriptase [Bio-Rad] using random and poly(A) priming. qRT-PCR was performed with KAPA SYBR fast [KAPA Biosystems] in 384-well format with a total reaction volume of 10µl. For measuring the expression levels of GAPDH and OCT4, cDNA was diluted 1:10 and 2µl were used for qPCR. Relative expression levels were calculated based on Δ/ΔCt and/or ΔCt analysis. qRT-PCR primers used in this study were:

GAPDH fw (CAGTCTTCTGGGTGGCAGTGA), GAPDH rev (CGTGGAAGGACTCATGACCA), OCT4 fw (CGTTGTGCATAGTCGCTGCT), OCT4 rev (GCTCGAGAAGGATGTGGTCC),

### sgRNAs

TPP1S KO: gtgtagccgtggggatggca; TPP1L KO: tgacgaacggcccaaatgcc; TPP1^L104A^: ctggattcgggagctgattc, TPP1 KO: ccctgatacgtccgacgtcg. The TPP1 KO strategy to disrupt both isoforms has been independently validated in the HCT116 cancer cell line (Vogan *et. al*, in preparation).

### IF/TIF analysis

For analysis by IF, cells were washed with PBS, fixed with 3.7% formaldehyde in PBS, permeabilized and blocked with PBS + 1mg/mL BSA, 3% v/v horse serum, 0.1% Triton X-100, and 1mM EDTA. Fixed cells were incubated with antibodies against TRF1 raised in rabbit against amino acid residues 17-41 (856-R1), and γ-H2AX (Millipore) in blocking solution followed by secondary antibodies. Scoring of TIF positive cells was performed single blind. Box plots show scoring of 9-11 fields of view and present median, mean, 95% confidence interval and outliers. *p*-values were determined using Prism 7’s Mann-Whitney test.

### Metaphase Spreads/FISH

Cells were treated with colcemid at 100μg/ml for 2 hours. We collected the cells using trypsin and incubated at 37C in prewarmed 75mM KCl. The cells were spun down and the KCl was removed. Cells were slowly resuspended in a fixative of 3:1 methanol:acetic acid. Cells were stored over night at 4C. The next day cells were spread, dropwise, onto microscope slides and washed twice with 1m 3:1 methanol:acetic acid solution. Slides were then placed onto a 70ºC humidified heat block for 1 minute.

Telomeres were detected by a previously described protocol (Lansdorp, Verwoerd et al., 1996) with a few minor changes. We fixed slides in 4% PFA diluted in PBS then treated with pepsin (1mg/ml) prepared in 10mM glycine pH 2 and warmed up to 37 degrees. After one more wash in 4% PFA, slides were washed with PBS, then dehydrated in an ethanol series. Each slide received 70 µl of hybridization mixture and was denatured at 65 degrees for 5 minutes, and then hybridized overnight with a Cy3 Tel-C PNA probe (PNA Bio) at room temperature in a hybridization chamber. The next day the slides were washed with 70% formamide, 10mM Tris-HCl pH 7.2, 0.1% BSA solution, and then with 0.1M Tris-HCl pH2, 0.15M NaCl, 0.08% Tween, with DAPI (diluted 1:1000 from 10mg/ml stock) added to the second wash. Coverslips were mounted with ProLong^®^ Gold Antifade Mountant (Thermo Fisher Scientific. All microscopy (IF and metaphase spreads) were imaged on a Nikon Eclipse TE2000-E epifluorescent microscope equipped with an Andor Zyla sCMOS camera.

## Acknowledgements

We would like to thank the Collins and Hockemeyer lab members for helpful discussion. Kunitoshi Chiba for technical assistance with TRAP, Stephen Floor for TrIP-Seq data, and Brendan Finnerty for blind scoring of images. D.H. is a Pew-Stewart Scholar for Cancer Research supported by the Pew Charitable Trusts and the Alexander and Margaret Stewart Trust. The work in the Hockemeyer laboratory is supported by the Siebel Stem Cell Institute and NIH R01-CA196884. Work in the Collins lab was supported by NIH R01-HL079585.

## Author contributions

JMB and DH conceived of all the experiments. JMB conducted all the experiments related to TPP1 L104A and isoform knockouts. SGR conducted all the experiments related to TPP1^∆L^. KMH conducted all the experiments related to TPP1 knockout. JMV and XZ conceived of and validated the knock out strategy for TPP1 and provided the associated targeting plasmids. JMV and XZ conceived of, validated and provided plasmids for TPP1 overexpression from transgenes integrated at the *AAVS1* locus. DH and KC supervised research. JMB and DH wrote the manuscript with the assistance of the other authors.

---

**Figure EV1.**
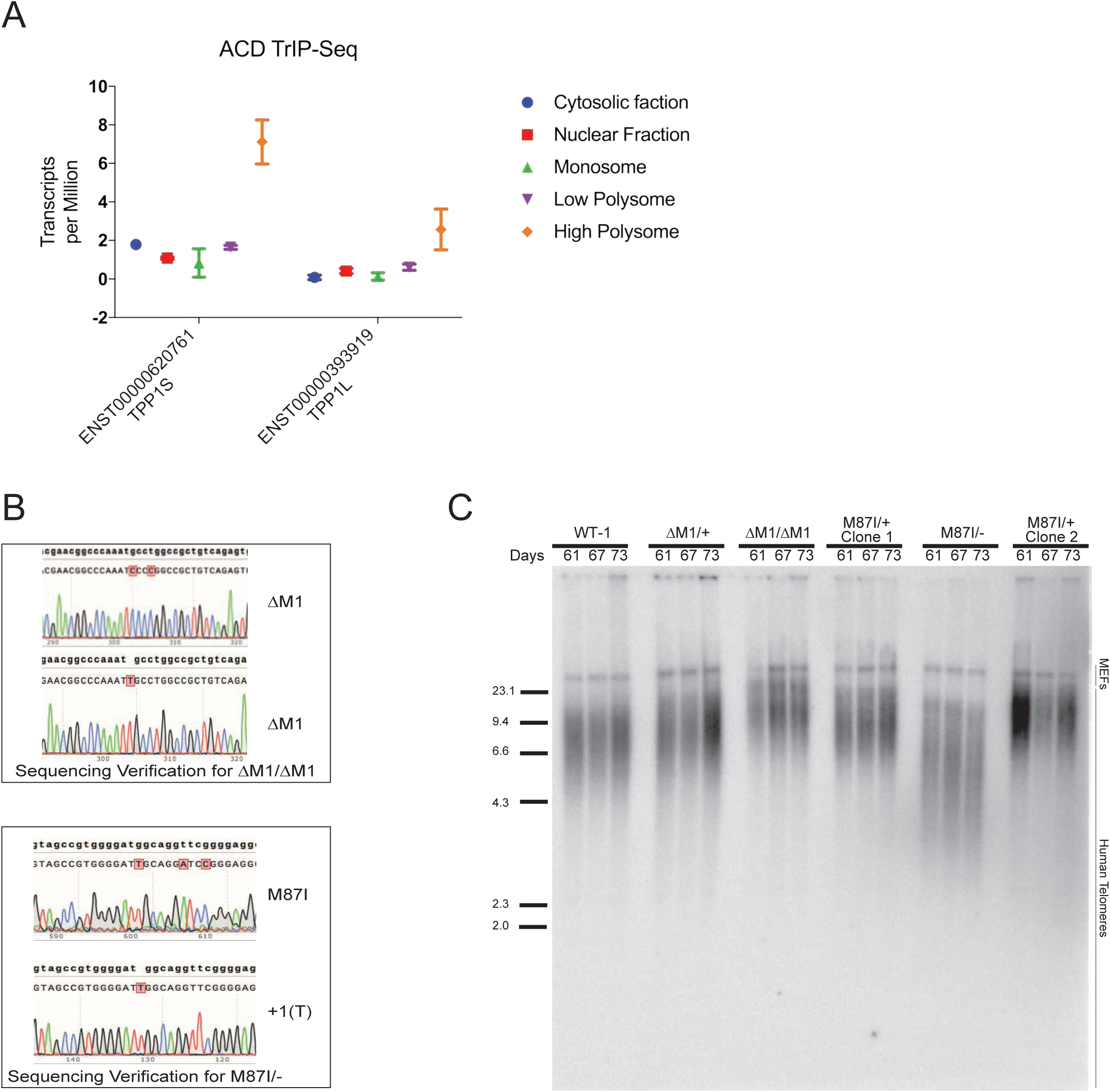
Confirmation of TPP1 isoform expression and ATG mutations. **A.** Transcript Isoforms in Polysomes sequencing (TrIP-seq) data for TPP1 isoforms from recent genome wide TrIP-Seq analysis. Nuclear and cytosolic fraction show total transcripts of each isoform per million transcripts indicating that TPP1S mRNA is more abundant than the mRNA of TPP1L. Not all transcripts are engaged by the ribosomes. The monosome represents isoform transcript that are associated with one ribosome, these tend to be enriched for retained introns and nonsense mediated decay targets compared to the polysome-bound fractions. The low polysome fraction represents isoform transcripts associated with low translation levels. The high polysome fraction represents isoforms that are highly translated. **B.** Sanger sequencing verification of mutations at the ATG’s of TPP1L and TPP1S KO cell lines. **C.** TRF analysis of additional TPP1 isoform KO clones. Heterozygous clones do not show a telomere length defect. Only TPP1^M87I/−^ showed progressive shortening.

**Figure EV2.**
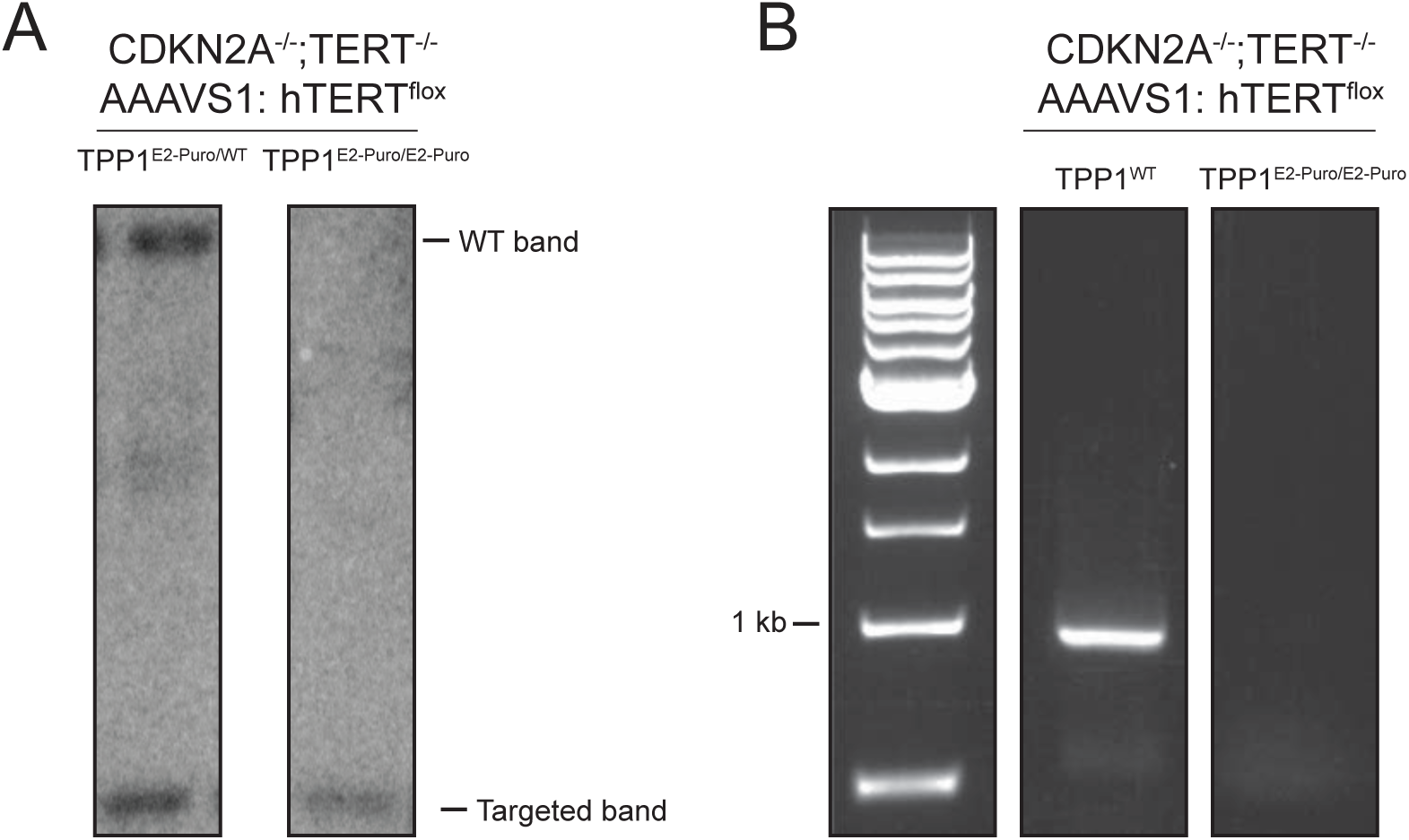
Confirmation of complete TPP1 knockout. **A.** Southern blot analysis of TPP1 locus in CDKN2A^−/−^, TERT^−/−^, AAVS1:TERT^flox^ and CDKN2A^−/−^, TERT^−/−^, TPP1^E2-Puro/E2-Puro^, AAVS1:TERT^flox^ cell lines showing insertion of puromycin cassette into exon 2 for a heterozygous and homozygous targeted clone. **B.** PCR genotyping analysis of clones shown in (A) demonstrating loss of the band for homozygously targeted clone.

**Figure EV3.**
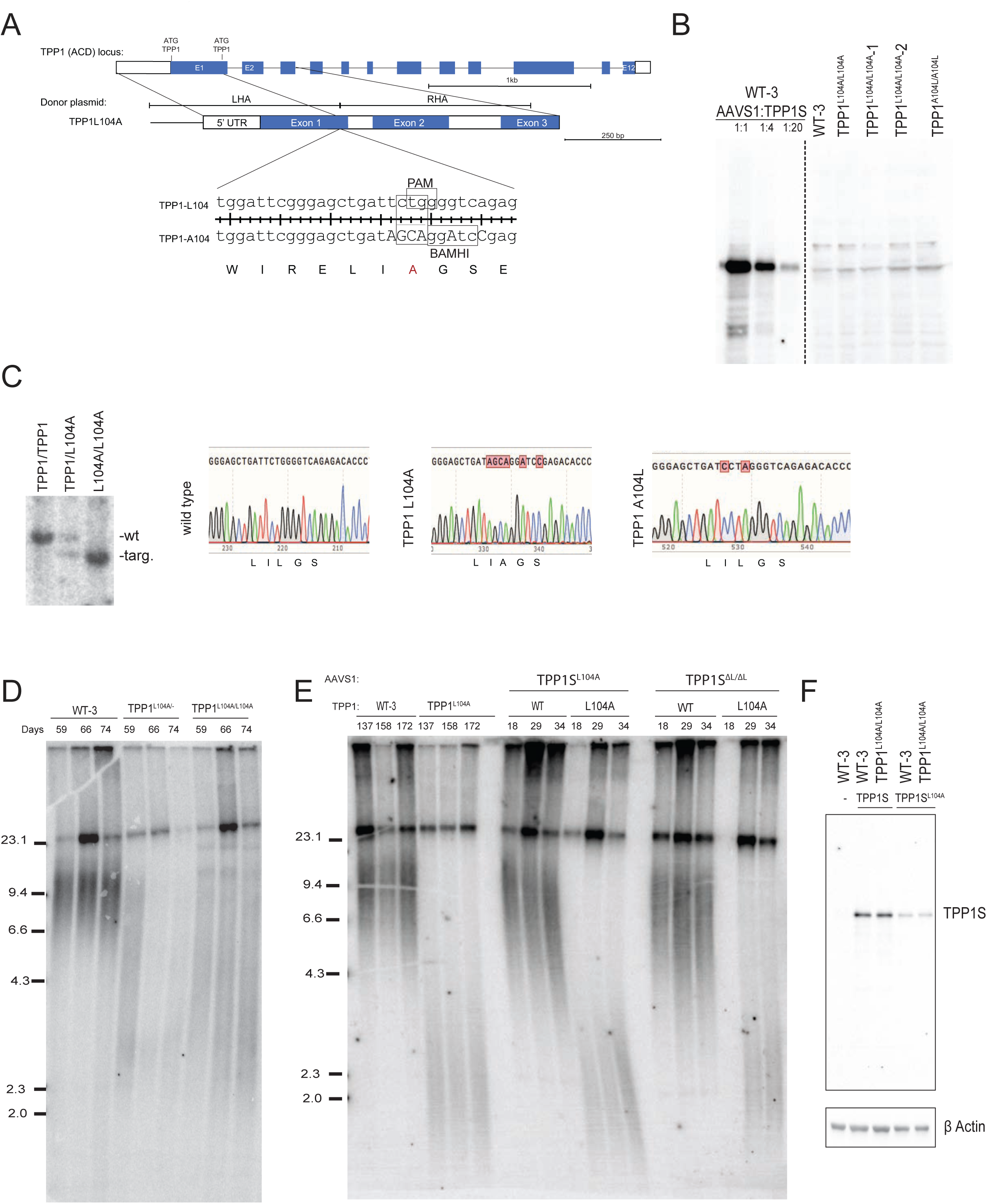
Analysis of TPP1 L104A and TEL patch deficient mutants. **A.** Targeting schematic for the generation of TPP1^L104A/L104A^. A donor plasmid carrying the L104A substitution and silent BamHI, to allow for genotyping, was used to modify the endogenous TPP1 locus. **B.** Left side, Western blot titration analysis of protein extracts taken from wild type cells overexpressing TPP1S. Right side, Western blot analysis of wild type, TPP1^L104A^, and TPP1^A104L^ repair clones. Samples collected 60 days post targeting repair of ACD locus. Parental samples were collected on the same day. **C.** Southern blot analysis of TPP1 L104A clones. Banding pattern indicates both heterozygous and homozygous knock in of mutant allele. Southern blot confirmed by Sanger sequencing. **D.** TRF analysis monitoring telomere length changes of early time points for TPP1^L104A/+^ and TPP1^L104A/L104A^ hESCs. Numbers indicate days post targeting. **E.** TRF analysis of telomere length changes in TPP1^L104A/L104A^ and wild type cell lines following targeting the *AAVS1* locus with either TPP1S^L104A^ or TPP1S^∆L^ AAVS1 overexpression constructs. Numbers indicate days after targeting. Parental cell lines samples were collected on the same days as overexpression samples. **F.** Western blot analysis of TPP1 overexpression for cell lines in D. Samples were collected 32 days post electroporation of overexpression cassettes.

**Figure EV4.**
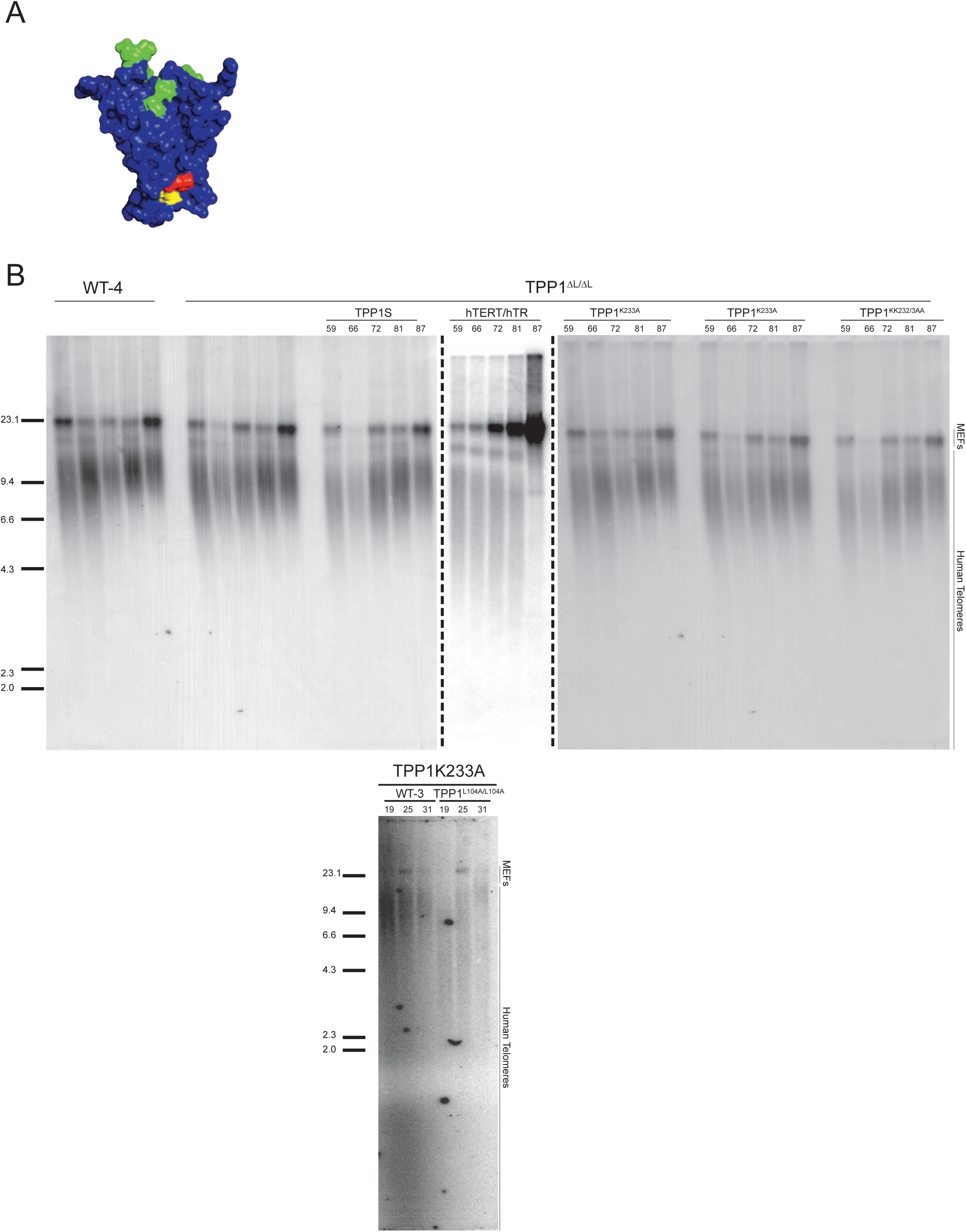
Extended TRF of TRF1 dominant negative expression in TPP1^L104A^ and control. Complementation analysis of TPP1^L104A^ by overexpression of dominant negative allele of TRF1, TRF1^∆A∆M^. Numbers indicate days post targeting TPP1^L104A/L104A^ hESCs in the *AAVS1* locus with a TRF1^∆A∆M^ overexpression cassette.

**Figure EV5.**
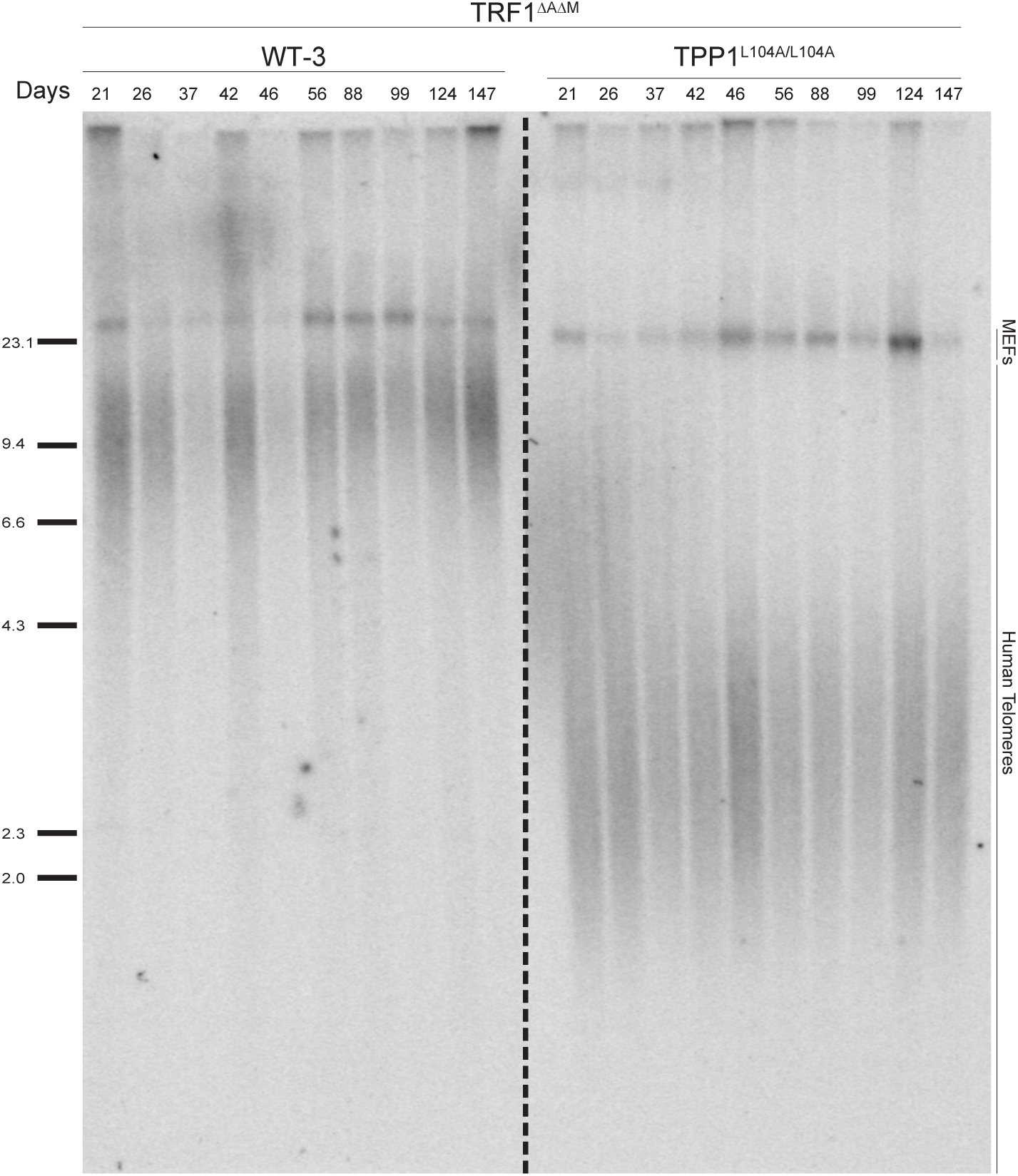
TPP1^∆L/∆L^ and ubiquitin deficient mutations. **A.** Space filling model of the TPP1 OB fold (Wang et al., 2007). The TEL patch is shown in green, residue L104 is shown in read and residue K233 is shown in yellow. **B.** Complementation analysis of TPP1^∆L/∆L^ and TPP1^L104A/L104A^ cell lines with TPP1 mutants incapable of being ubiquitinated at K232, K233 or both. TPP1^∆L/∆L^ is not responsive to hTERT/hTR overexpression. Numbers indicate days post-electroporation of TPP1 repair cassettes.

**Table EV1:** Cell lines generated. All cell lines were generated in the WIBR#3 hESC line (NIH stem cell registry #0079). Wild type cell lines are subclones of WIBR#3 isolated and cultured in parallel with their respective experimental line. TPP1^∆L/∆L^ parental line was previously published (Sexton et al., 2014).

| Genotype (all in WIBR#3, NIH line # 0079) | Cell line | Figure introduced |
| --- | --- | --- |
| TPP1<sup>M1Δ/M1Δ</sup> | JMB1 | Fig. 1 |
| TPP1<sup>M87I/-</sup> | JMB2 | Fig. 1 |
| CDKN2A<sup>-/-</sup>, TERT<sup>-/-</sup>, TPP1<sup>wt/wt</sup>, AAVS1:F-TERT<sup>flox</sup> | SR1 | Fig. 3 |
| CDKN2A<sup>-/-</sup>, TERT<sup>-/-</sup>, TPP1<sup>E2-Puro/E2-Puro</sup>, AAVS1:F-TERT<sup>flox</sup> | KMH1 | Fig. 3 |
| CDKN2A<sup>-/-</sup>, TERT<sup>-/-</sup>, TPP1<sup>E2-Puro/wt</sup>, AAVS1:F-TERT<sup>flox</sup> | KMH2 | EV Fig. 2 |
| TPP1<sup>L104A/L104A</sup> | JMB3 | Fig. 4 |
| TPP1<sup>A104L/A104L</sup> | JMB4 | Fig. 4 |
| TPP1<sup>L104A/L104A</sup>-1 | JMB5 | Fig. 4 |
| TPP1<sup>L104A/L104A</sup>-2 | JMB6 | Fig. 4 |
| WT-3; AAVS1:F-ΔPBD | JMB7 | Fig. 4 |
| TPP1<sup>L104A/L104A</sup>, AAVS1:F-ΔPBD | JMB8 | Fig. 4 |
| TPP1<sup>M87I/-</sup>; AAVS1:myc-POT1ΔOB | JMB9 | Fig. 6 |
| TPP1<sup>M87I/-</sup>; AAVS1:F-hTERT/hTR | JMB10 | Fig. 6 |
| WT-3; AAVS1:F-TRF1 | JMB11 | Fig. 6 |
| TPP1<sup>L104A/L104A</sup>, AAVS1:F-TRF1 | JMB12 | Fig. 6 |
| WT-3; AAVS1:TRF1<sup>ΔAΔM</sup> | JMB13 | Fig. 6 |
| TPP1<sup>L104A/L104A</sup>, AAVS1:F-TRF1<sup>ΔAΔM</sup> | JMB14 | Fig. 6 |
| WT-3; AAVS1:F-GFP | JMB15 | EV Fig. 1 |
| TPP1<sup>L104A/L104A</sup>, AAVS1:F-GFP | JMB17 | EV Fig. 1 |
| TPP1<sup>ΔM1/+</sup> | JMB18 | EV Fig. 1 |
| TPP1<sup>M87I/+</sup> Clone 1 | JMB19 | Fig. 1 |
| TPP1<sup>M87I/+</sup> Clone 2 | JMB20 | Fig. 1 |
| TPP1<sup>+/L104A</sup> | JMB21 | EV Fig. 3 |
| TPP1ΔL/ΔL; AAVS1:3xF-TPP1S | SR2 | EV Fig. 3 |
| TPP1ΔL/ΔL; AAVS1: F- hTERT/hTR | SR3 | EV Fig. 3 |
| TPP1ΔL/ΔL; AAVS1: 3xF-TPP1S<sup>K232A</sup> | SR4 | EV Fig. 3 |
| TPP1ΔL/ΔL; AAVS1:3xF-TPP1S<sup>K233A</sup> | SR5 | EV Fig. 3 |
| TPP1ΔL/ΔL; AAVS1:3xF-TPP1S<sup>KK232/3AA</sup> | SR6 | EV Fig. 3 |
| WT-3; AAVS1:3xF-TPP1<sup>K233A</sup> | JMB22 | EV Fig. 3 |
| TPP1<sup>L104A/L104A</sup>, AAVS1:3xF-TPP1<sup>K233A</sup> | JMB23 | EV Fig. 3 |

